# Trifluoperazine exhibits broad-spectrum antiviral activity against arboviruses

**DOI:** 10.64898/2026.03.17.712523

**Authors:** Laxmi Mishra, Manjula Kalia

## Abstract

The recurrent outbreaks and geographical expansion of mosquito-borne arboviruses pose a significant challenge to public health worldwide. The disease outcome for arboviral infections ranges from acute febrile illness to severe conditions such as encephalitis, hemorrhagic shock, and mortality. Current treatment options for these viruses are limited to supportive care, necessitating an urgent need for a safe and effective broad-spectrum antiviral. In this study, we have identified Trifluoperazine (TFP), an FDA-approved antipsychotic, as a potent broad-spectrum antiviral against Japanese encephalitis Virus (JEV), Dengue virus (DENV) and Chikungunya virus (CHIKV) infections. The antiviral effect of TFP was also seen in the animal models of JEV and CHIKV with significantly reduced disease severity. Mechanistically, TFP treatment increased the phosphorylation of eIF2a and induced an adaptive ER stress response in diverse cell types. Alleviation of TFP-induced ER stress by chemical chaperone 4PBA abolished the antiviral activity of the drug and rescued virus replication in cells. The robust *in vitro* and *in vivo* efficacy of the drug against arboviruses highlights the potential for repurposing TFP as a broad-spectrum antiviral candidate.

## 1. Introduction

Arboviruses, also known as arthropod-borne viruses, are RNA viruses primarily transmitted to humans by the bite of infected mosquitoes, ticks, and sand flies. They belong to families *Flaviviridae*, *Togaviridae*, *Bunyaviridae*, and *Reoviridae*. Earlier limited to tropical and sub-tropical regions, these viruses have now become major global health threats, causing over 700,000 deaths and hundreds of millions of infections annually [1]. Among these, approximately 390 million cases are caused by the Dengue Virus (DENV), with other major contributors being the Yellow Fever Virus (YFV), Chikungunya Virus (CHIKV), Zika Virus (ZIKV), Japanese Encephalitis Virus (JEV), and West Nile Virus (WNV). Together, these viruses put 70% of the world’s population at risk of infections [2].

Most of the arboviral diseases in Asia are caused by *flaviviruses* and *alphaviruses*. These include JEV, DENV and CHIKV [3]. The clinical manifestations associated with each of these arboviruses range from mild to severe symptoms. While mild symptoms include self-limited febrile-like conditions, the disease severity in JEV infection is characterized by encephalitis, in DENV infections by hemorrhagic shock, and in CHIKV infections by severe myalgia and arthralgia. There are no antivirals available and current treatment is restricted to symptomatic relief. The development of broad-spectrum antivirals against these viruses can not only reduce the burden on healthcare systems but also alleviate socioeconomic pressure caused by these viral infections.

Targeting the host can serve many advantages for broad-spectrum antiviral strategies. It includes a large number of highly conserved target repertoires, barrier to viral resistance [4], and moderation of detrimental host responses [5]. Additionally, host-directed antivirals can offer protection against future emerging viruses, a concern often associated with geographically expanding and re-emerging arboviruses. Hence, targeting host cell machinery, crucial and fundamental to many host-virus interactions, can be an effective antiviral approach.

The infection cycle of many arboviruses depends on the endoplasmic reticulum (ER). This complex organelle supports the various stages of virus replication, including the translation of the RNA genome and the establishment of the replication complex [6, 7]. As the infection progresses, the accumulation of viral proteins in the ER triggers the activation of intracellular stress responses [8–10]. It primarily includes ER-associated degradation (ERAD), autophagy, apoptosis, and inflammation, which are influenced by the unfolded protein response (UPR), also known as ER stress response [11]. The interplay of these pathways exerts antiviral effects on the life-cycle and propagation of many viruses [12–17]. Additionally, the induction of ER stress by small molecules has been shown to have an antiviral effect against a wide range of RNA viruses [18–21].

Several FDA-approved phenothiazines have been shown to modulate ER stress to exhibit anti-tumorigenic [22, 23], neuroprotective [24], and antiviral [25] activity. Recently, a screening of autophagy inducers in our lab had identified the ER stress-dependent anti-JEV activity of phenothiazine derivative, methotrimeprazine (MTP) [26]. This study had also shown robust antiviral effect of Trifluoperazine (TFP) in cell line and sub-lethal mice model. TFP subdued the JEV infection with high-potency at lower concentrations, suggesting that the therapeutic effects of TFP can be achieved at low and clinically favorable doses. Also, TFP has been shown to target distinct molecular mechanisms essential for different viruses [27–30]. The well-established pharmacological profiles, high potency and specific modulation of conserved host pathways encouraged us to further explore the repurposing potential of TFP as a broad-spectrum, host-directed antiviral candidate against arboviruses.

Here, we describe TFP to be a potent inhibitor of JEV, DENV-2, and CHIKV infections. TFP induced ER stress in diverse cell types, and pharmacological rescue of TFP-induced ER stress reverses the antiviral effect of the drug, suggesting its the broad-spectrum antiviral activity is mediated through ER stress. The antiviral efficacy of the drug was also demonstrated in preclinical models of JEV and CHIKV infections. However, the lack of an immunocompetent animal model for DENV-2 limited the *in vivo* evaluation of TFP against dengue.

## 2. Materials and Methods

### 2.1. Ethics statement

All animal experiments were performed as per the CCSEA (Committee for Control and Supervision of Experiments on Animals) guidelines, mandated by Government of India. The Institutional Animal Ethics Committee of the Regional Centre for Biotechnology reviewed, approved, and monitored animal experiments under protocol number RCB/IAEC/2023/168 and RCB/IAEC/2025/225. All infection experiments were performed in Biosafety Laboratory 2 (BSL2) facility.

### 2.2. Cell lines and Viruses

Neuro2a (mouse neuroblastoma), Huh7 (human hepatocarcinoma), Vero (monkey kidney epithelial cells), C6/36 (mosquito cell line) and BHK-21 (baby hamster kidney cells) were obtained from the cell repository at National Centre for Cell Sciences (NCCS) Pune, India. ERMS (human embryonal rhabdomyosarcoma) cell line was obtained from American Type Cell Collection (ATCC, USA) (RD-CCL-136-ATCC). All cell lines were mycoplasma negative. Neuro2a, Huh7, ERMS and L929 were cultured in Dulbecco’s modified Eagle’s Medium (DMEM; Gibco 12100-046). Vero, C6/36 and BHK-21 were cultured in Eagle’s minimum essential medium (MEM; Gibco 61100-061). Both media were additionally supplemented with 10% fetal bovine serum (FBS; Gibco 10270-106), 100 µg/ml penicillin-streptomycin (PS; Himedia A007-100ML), and 2 mM L-glutamine (G; Himedia TCL012). C6/36 cells were grown at 28°C with 5% CO2, and all mammalian cell lines were grown at 37°C with 5% CO2.

JEV isolate Vellore P20778 strain (GenBank accession no. AF080251) and DENV-2 isolate strain P23085 INDI-60 (GenBank accession no. KJ918750) were propagated in C6/36 cell line. CHIKV isolate IND-06-Guj (GenBank accession no. JF274082) was generated in BHK-21. This cell-cultured CHIKV was used for both, *in vitro* and *in vivo,* experiments. For animal experiments of JEV, the mouse-adapted JEV S3 isolate were propagated in 2-3 days old pups [31]. For animal experiment of DENV, mouse-adapted DENV-2 (P8-P23085 INDI-60) was propagated in C6/36 cells [32]. All the virus strains were titrated in Vero cells using plaque assays and foci forming assays.

### 2.3. Isolation and culture of bone marrow-derived macrophages (BMDMs)

BMDMs were isolated from 6-8 weeks old C57BL/6 or AG129 mice. These mice were euthanized and femurs were dissected out. After washing with PBS and RPMI media (Gibco 31800-022), bone marrow was flushed out using syringe needle. Extruded bone marrow cells were collected in RPMI, supplemented with 10% L929-conditioned media. L929-conditioned media is cell culture supernatant, rich in Macrophage Colony Stimulating Factor (M-CSF), primarily used to differentiate bone marrow cells into macrophages. Conditioned media was prepared by starving L929 cells in DMEM (10% FBS, 1X PSG) for seven days. The collected bone marrow cells were subjected to RBC lysis (RBC lysis buffer; GCC Biotech 19114B1076) and subsequently grown in RPMI supplemented with L929-conditioned media for 7 days. On 7^th^ day, BMDMs were detached using 10mM EDTA (Sigma E9884) and seeded in 24-well plates to perform antiviral assays.

### 2.4. Cytotoxicity assay

To determine the cytotoxicity of TFP, cells were seeded in 96-well plates. After preculturing for 16 hours, cells were treated with media containing different concentrations of TFP/DMSO. Drug incubation time for Neuro2a and ERMS was 24 h, and for Huh7 was 72 h. To determine cell viability, MTT assay (VWR life science 0793) was performed as per the manual. The percent cell viability was calculated and plotted as: [(Absorbance for test condition/Absorbance for DMSO treated condition) X 100].

### 2.5. Virus infection and cell treatment

Based on the tropism of these viruses, distinct cell lines were used to test and characterize the antiviral potency of the drug. Neuro2a cells were used for JEV infections, Huh7 were used for DENV infections and ERMS were used for CHIKV infections. All these cells were incubated with virus for 1 h (JEV and CHIKV) or 2 h (DENV-2) at 37°C. Post-incubation, cells were washed with PBS and complete media with/without drug was added. Control cells were treated with DMSO and test groups were treated with TFP till the time of harvest. All JEV and CHIKV experiments were performed with 1 MOI, and DENV-2 experiments were performed with 2 MOI. Upon completion of experiment, cell culture supernatants were collected to quantify virus titers through plaque/foci forming assays, and cells were harvested for qRT-PCR. To study ER stress, cells were treated with DMSO/Tg (1 µM)/TFP (10 µM)/4PBA (2 mM)/TFP+4PBA for 3 h and 6 h, and cell lysates were processed for western blotting. All experiments had biological triplicates and were repeated twice or thrice. Tg (T9033), TFP (T8516), and 4PBA (1716-12-7) were purchased from Sigma Aldrich.

### 2.6. RNA isolation and quantitative real time PCR (qRT-PCR)

Total RNA was isolated with phenol-chloroform separation method using RNAiso Plus (Takara 9109). cDNA was prepared from 1 µg RNA, using random hexamers (Sigma H0268) and ImProm-II^TM^ Reverse Transcription System kit (Promega A3800). qRT-PCR was set up on QuantStudio 6 flex. For qRT-PCR, SyBr mix (Takara RR420A) was used to determine the expression of *GAPDH*, DENV and CHIKV RNA, and TaqMan mix (Takara RR390A) (with specific probe) was used for JEV RNA. Here *GAPDH* was used as a housekeeping control. qRT-PCR for each biological sample was performed in technical duplicates. The sequences of primers and probe used in this study are as follows (5’-3’): Human GAPDH: F-TGCACCACCAACTGCTTAGC, R-GGCATGGACTGTGGTCATGAG; Mouse GAPDH: F- CGTCCCGTAGACAAAATGGT, R- TTGATGGCAACAATCTCCAC; JEV: F-AGAGCACCAAGGGAATGAAATAGT, R- AATAAGTTGTAGTTGGGCACTCTG; DENV-2: F1-GAGAGACCAGAGATCCTGCTGTCT, F2- GAAAGACCAGAGATCCTGCTGTCT, R-ACCATTCCATTTTCTGGCGTT; CHIKV: F-GGCAGTGGTCCCAGATAATTCAAG, R-GCTGTCTAGATCCACCCCATACATG; and JEV probe: CCACGCCACTCGACCCATAGACTG with 5’ 6-FAM as reporter/3’ TAMRA as quencher.

### 2.7. Western Blotting

Lysates were prepared by incubating cells in lysis buffer (150 mM NaCl, 1% Triton X-100, 50 mM Tris-HCl pH 7.5, 1 mM PMSF (Sigma 329-98-6), and protease inhibitor cocktail (Sigma (P8340)) for 1.5 h at 4°C. BCA (Bicinchoninic Acid) assay (G-Biosciences 786-570) was performed to estimate the protein concentration in cell lysates. 5X loading buffer (40% glycerol, 20% β-mercaptoethanol, 0.04% bromophenol blue, 6% SDS, 0.25M Tris-HCl pH 6.8) was added to the lysates and heated at 95°C for 10 minutes to allow denaturation of proteins. Equal amounts of protein were loaded and separated through SDS-PAGE. Proteins were transferred from gel to polyvinylidene fluoride (PVDF) membrane (Merck Millipore IPVH00010). After the complete transfer, blocking was done using 5% skimmed milk in 1X PBS (1 h at room temperature). Primary antibodies were prepared in 5% BSA in 1X PBS and added onto the blots for overnight at 4°C. Membrane was washed thrice with 1X PBST (0.1% Tween20 in 1X PBS) and then incubated with HRP-conjugated secondary antibodies, diluted in 5% milk in 1X PBS, for 1 h at RT. After three more washes with 1X PBST, blots were developed using HRP substrate. Visualization was done on Gel Doc XR+ gel documentation system (Bio-Rad). ImageJ (NIH, USA) software was used to quantify the intensity of bands. Fold change was calculated by normalizing with loading control. The following primary antibodies were used in the study: p-eIF2a (ab32157), eIF2a (ab5369), PERK (CST#5683), and GAPDH (GTX100118). Horseradish peroxidase (HRP)-conjugated secondary antibodies were purchased from Jackson Immunochemicals.

### 2.8. Plaque and Foci-forming assays

Titers of JEV and CHIKV were calculated using plaque assay, and of DENV-2 using foci-forming assay. Vero cells were infected with serially diluted virus supernatant. The incubation time with infection media was 2 h for DENV-2 and 1 h for JEV and CHIKV. Post-incubation, cells were washed with 1X PBS and media overlay was added. For plaque assay, 2% low-melting agarose (Sigma A4018) was mixed with 2X MEM in 1:1 ratio. Once agarose plug was solidified, the plates were kept in incubator at 37°C. JEV infected plaque plates were incubated for 5.5 days and CHIKV infected plates were incubated for 36-48 hours. After incubation, cells were fixed with 3.7% formaldehyde for 4 h, agarose plug was removed and the fixed cells were stained with 0.1% crystal violet. For foci forming assay, only MEM media was added as overlay and infected cells were incubated for 72 h. After the incubation period was over, cells were washed and fixed with 2% para-formaldehyde. Fixed cells were immunostained with 4G2 primary antibody. Alexa Fluor 488 labeled secondary antibody was used and foci was observed under fluorescence microscope. Number of plaque forming units (pfu) and foci forming units (ffu) were counted to calculate the titer. The pan-flavi antibody 4G2 was harvested from HB-112 (ATCC) culture supernatant and Alexa 488 fluorophore-coupled secondary antibodies were purchased from Invitrogen, Thermo Fisher Scientific.

### 2.9. Animal experiments

All mice were housed in well-ventilated cages with light/dark cycle of 12 h/12 h, and temperature and humidity of 20-24°C and 30-70%, respectively. Food and water supply were provided *ad libitum*. Blinding protocol was not used for any of the following animal experiments.

#### 2.9.1. JEV

3 weeks old C57BL/6 mice of either sex were grouped into mock/JEV/JEV+TFP, without any bias. For survival assay, these mice were intraperitoneally injected with either JEV S3 isolate (10^7 pfu/mice) or equal volumes of incomplete DMEM. At 4 hpi, mice were orally treated with 100 µl of vehicle (PEG 400; Sigma 202398) or TFP (1 mg/kg). Drug treatment was given everyday till 8 days post infection (dpi). All mice were weighed till 10 dpi. During this time, mice were also monitored for other symptoms of JEV such as piloerection, hunchback, hind-limb paralysis and mortality. Survival curve was plotted and analysis was done.

To determine the viral load in brain, 3 mice from each group (mock/JEV/JEV+TFP) were sacrificed at 5, 6 and 7 dpi. Mice brain was collected and homogenized in incomplete DMEM. Tissue homogenate was centrifuged and supernatant was used to perform plaque assay.

#### 2.9.2. DENV

For DENV animal experiments, 6-8 weeks old AG129 mice (IFN α/β/γR-/- 129/Sv) of either sex were used. These mice were randomly divided into mock/DENV/DENV+TFP groups. Mice were infected with 100 µl of mouse-adapted DENV-2 (10^5 ffu/mice), intraperitoneally or inoculated with equal volumes of incomplete DMEM. At 4 hpi, drug dosing began and mice were treated with vehicle (PEG 400) or TFP (1 mg/kg/day) for 8 days. Infected and drug-treated mice were monitored for weight loss, symptoms like hunch back and ruffled fur, and mortality.

#### 2.9.3. CHIKV

8-10 weeks old C57BL/6 mice of either sex were divided into three groups- Mock, CHIKV, CHIKV+TFP. Mock mice were injected with PBS and mice from other two groups were infected with 10^4 pfu of CHIKV (INDI-06-Guj strain). 50 µl of infection inoculums was injected subcutaneously at the ventral side of each hind foot. First dose of TFP (1 mg/kg) was given at 4 hpi, with subsequent doses at an interval of 24 hours till 8 dpi. Route of administration for drug was oral. Mice were followed up till day 10 and parameters such as body weight and paw edema were recorded. To measure the edema in foot, a digital plethysmometer was used and readings were documented. On day 2 and day 3 post-infection, the retro-orbital blood was collected to quantify the viral load in serum.

### 2.10. Evans blue leakage assay

To quantify the blood brain barrier (BBB) breach in mice, 100 µl of 2% Evans blue dye (Sigma E2129-10G) was injected intraperitoneally. After 1.5 h mice were sacrificed, and brains were carefully harvested. Tissue homogenate was prepared by homogenizing the brain in dimethylfumarate (DMF; Sigma 242926-25G) (500 µl DMF for 200 mg tissue). Dye was extracted from tissue by heating the homogenate overnight at 60°C. These samples were centrifuged and supernatant was used to measure the absorbance of dye at 620 nm. A range of different concentrations of Evans blue were used to plot the standard curve and the dye’s concentration in brain samples was calculated.

### 2.11. Cytokine Bead Array (CBA)

CBA was performed to measure the cytokine levels in JEV infected mice brain. 3 mice were sacrificed from each group (mock/JEV/JEV+TFP) at 5, 6 and 7 dpi. Brain tissues were collected and lysates were prepared by homogenizing the tissue in lysis buffer. BCA was used to determine the concentration and 30 µg protein of each sample was used to quantify the cytokines. Assay kit of LEGENDPLEX MU (Biolegend 740622) Mix and Match panel of following cytokines- IL6, TNFa, RANTES and IFNB1 was used. For quantification, LEGENDplex^TM^ Multiplex Assay Software Qognit was used in which the concentration of cytokines were determined using standard curves.

### 2.12. Statistical Analysis

CC50 and IC50 values were calculated using dose-response curves. Statistic analysis was performed using unpaired Student’s t-test, one-way ANOVA followed by Dunnett test, and Mantel-Cox Log-rank test. Data significance was considered at p-value <0.05. Error bar indicates mean ± SD/SEM of data. All the graphs were plotted and analyzed using GrpahPad prism 8.

## 3. Results

### 3.1. Trifluoperazine exhibits broad-spectrum antiviral activity against RNA viruses

Previously, we have reported the antiviral potential of antipsychotic drug TFP against JEV [26]. In this study we explored the broad spectrum antiviral activity of TFP against three clinically relevant arboviruses- JEV, DENV and CHIKV. We assessed the antiviral activity of TFP by measuring the intracellular viral RNA and extracellular infectious virus particles of JEV, DENV-2 and CHIKV in untreated and drug-treated conditions (Fig. 1A-F). We observed a time dependent enhancement in viral RNA and titers, which confirmed efficient replication and propagation of these viruses in the respectively chosen cell line models. Conversely, treatment with TFP showed a robust reduction in the replication of these viruses. We observed 50-90% reduction in JEV (Fig.1A), DENV-2 (Fig.1C) and CHIKV (Fig.1E) RNA levels, and 50-70% reduction in their respective titers (Fig.1B, D, F). As observed for different time points, the antiviral activity of TFP against these RNA viruses is consistent and profound.

**Fig. 1:**
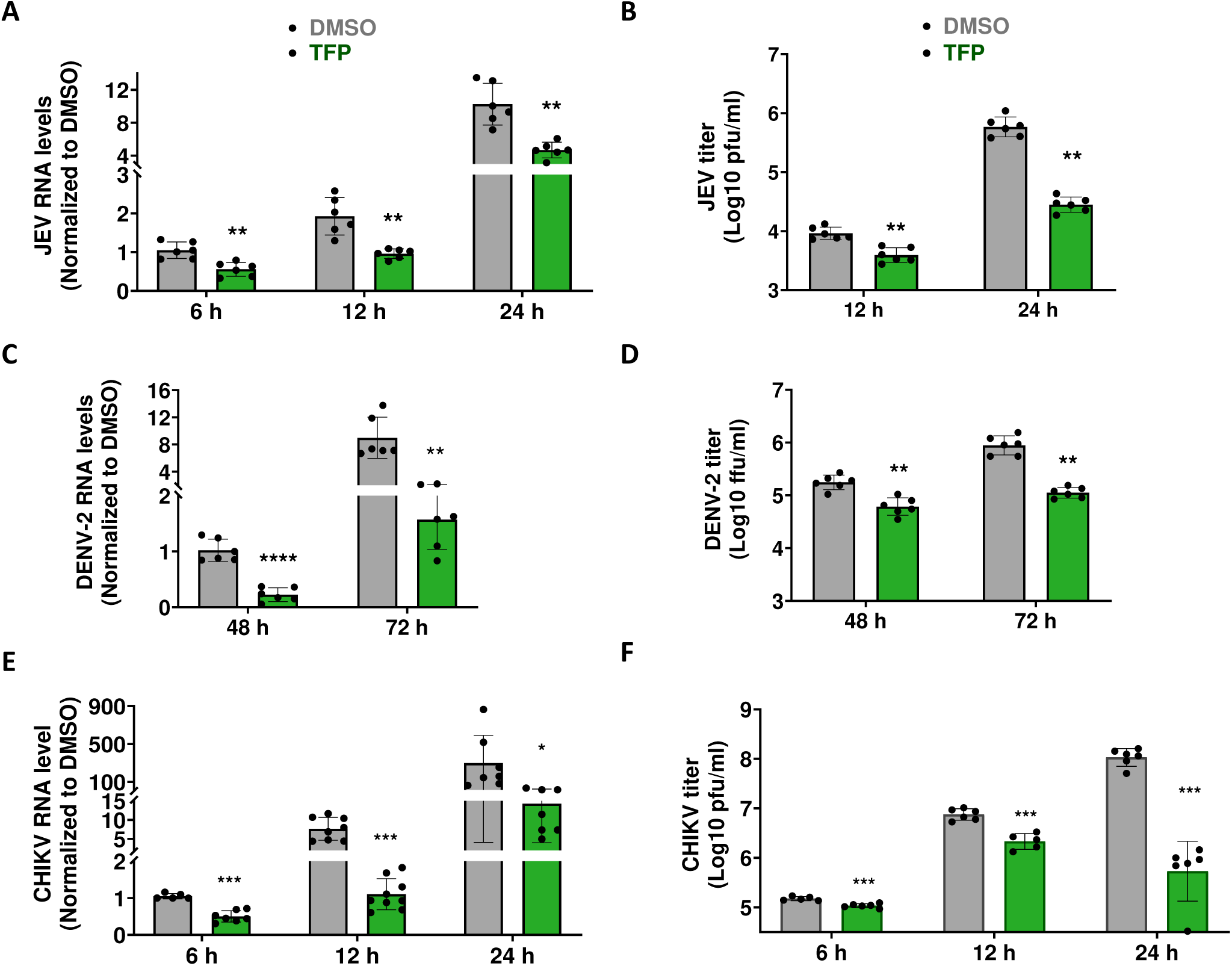
TFP shows broad-spectrum antiviral activity against JEV, DENV and CHIKV. (A, B) Neuro2a cells were infected with JEV at 1 MOI. After 1 h of incubation, cell culture medium was supplemented with DMSO (control) or TFP (10 µM). Cells were harvested at 6, 12, and 24 h post-infection (hpi). (A) qRT-PCR was performed to quantify JEV RNA levels, where data were normalized to DMSO at 6 hpi. (B) Culture supernatant was used to titrate the JEV using the plaque assay. (C, D) Huh7 cells were infected with DENV-2 at 2 MOI and treated with DMSO or 10 µM TFP for the indicated time points. (C) Intracellular DENV-2 RNA was quantified using qRT-PCR, and (D) the virus titers were determined by the foci-forming assay. The DENV-2 RNA was normalized to the 48 h DMSO control. (E, F) ERMS cells were infected with CHIKV at 1 MOI and treated with DMSO or TFP (10 µM) for the indicated time points. (E) Cells were harvested to quantify intracellular CHIKV RNA levels, and (F) culture supernatant was collected to determine the CHIKV titer by plaque assays. The viral RNA levels were normalized to 6 h DMSO control. Values were plotted from two independent experiments (*n*=6). The unpaired Student’s t-test was used to calculate the *p* values: \**p*<0.05, \*\**p*<0.01, \*\*\**p*<0.001, \*\*\*\**p*<0.0001.

### 3.2. TFP treatment attenuates the JEV infection and protects the mice from virus-induced pathogenesis

We first evaluated CC50 and IC50 values of TFP to determine the therapeutic window of the drug. In Neuro2a, TFP exhibited CC50 of 30.8 µM and IC50 of 0.1 µM (Fig. 2A), with remarkably high SI of 308. Treatment with TFP also showed a dose-responsive reduction in intracellular JEV RNA (Fig. 2B).

**Fig. 2:**
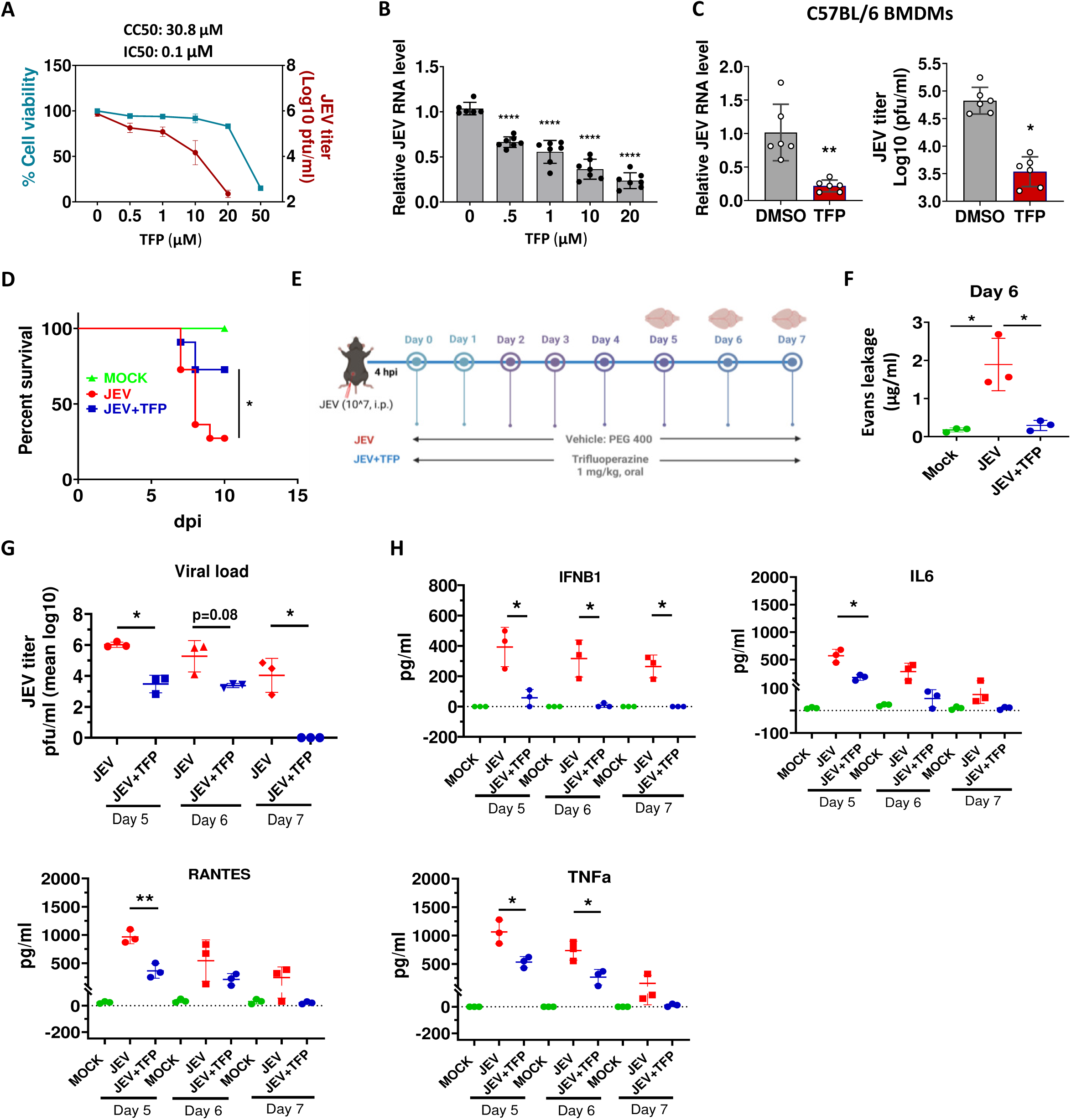
TFP effectively restricts JEV infection and virus-induced pathogenesis. (A, B) Neuro2a cells were mock/JEV-infected (1 MOI) and were treated with different concentrations of TFP until 24 hpi. (A) Culture supernatant was used to perform a plaque assay (*n*=6). Percent cell viability was measured using the MTT assay, and the data were normalized to the DMSO control (*n*=4). The graph represents the CC50 and IC50 values. (B) JEV RNA level was quantified and normalized to DMSO (n≥6). Statistical significance was determined using one-way ANOVA followed by Dunnett’s test. (C) BMDMs isolated from C57BL/6 mice were infected with JEV (5 MOI) and treated with 10 µM TFP for 24 h. qRT-PCR was performed to quantify the intracellular viral RNA, and a plaque assay was performed to quantify the virus titers. The relative levels of JEV RNA (normalized to DMSO) are shown in the left panel (*n*=6), and JEV titers are shown in the right panel (*n*=6). The unpaired Student’s t-test was used to determine the statistical significance. (D) 3-week-old C57BL/6 mice were mock-infected with DMEM or JEV-infected (10^7 pfu of JEV S3/mice), intraperitoneally. After 4 h, oral administration of vehicle (PEG 400) or TFP (1 mg/kg) was done at an interval of 24 hours for 8 days. Mice were monitored for encephalitis-like symptoms, and a survival curve was plotted for mock (*n*=4)/ JEV (*n*=11)/ JEV+TFP (*n*=11). Log-rank Mantel-Cox test was used to determine the statistical significance between vehicle-treated (JEV) and drug-treated groups (JEV+TFP). (E) Schematics of the experimental plan in mice to determine the effect of the drug on JEV-induced pathogenesis. C57BL/6 mice were JEV-infected and TFP-treated as mentioned above in the survival assay. (F) At day 6 p.i., mock or vehicle/TFP-treated JEV-infected mice (*n*=3 in each group) were inoculated with 2% Evans blue dye (100 µl) through an i.p. injection. These mice were euthanized, and brain tissue homogenates were prepared in DMF (200 mg/500 µl DMF). Absorbance of the dye was measured at 620 nm, and the concentration was determined from a standard curve. Statistical significance was determined using unpaired Student’s t-test. (G, H) Three C57BL/6 mice from each group of mock/JEV/JEV+TFP were sacrificed at days 5, 6 and 7 p.i. (G) Brain tissues were homogenized in DMEM, and supernatant was used to quantify viral load in brain using plaque assay. (H) Brain tissue lysates were prepared in cell lysis buffer, and an equal amount of protein (30 µg) from each sample was used to quantify cytokine levels using CBA kit. Data were analyzed using LEGENDplexTM Multiplex assay software. Each data point represents one mouse, and unpaired Student’s t-test was used to compare the statistical difference between JEV and JEV+TFP. \**p*<0.05, \*\**p*<0.01, \*\*\**p*<0.001.

Subsequently, we tested the efficacy of drug in *ex vivo* and *in vivo* system. We found that treatment with TFP significantly reduced the JEV infection in C57BL/6 derived BMDMs (Fig. 2C). Mice experiments from our previous study had shown that TFP improved the survival of mice infected with sub-lethal dose of JEV [26]. Here we evaluated the antiviral efficacy of drug in lethal model of JEV infected mice. Within 5-6 days of infection, we observed symptoms like significant loss in body weight with piloerection and hind-limb paralysis. Most of these symptomatic mice died within 2-3 days of symptom onset. Comparison of survival curves revealed that the proportion of mice survived in drug treated group, i.e. 72%, was significantly higher than that in the untreated group, i.e. 27% (Fig. 2D).

As JEV is a neurotropic virus, we next assessed the virus induced pathogenesis in brain of infected mice at 5, 6 and 7 dpi (experimental plan shown through schematics in Fig. 2E). JEV infected brains showed leakage of Evans blue dye on 6 dpi, which indicates permeability of BBB. However this breach was rescued in TFP treated group (Fig. 2F). We observed that viral titers in brain of drug treated mice had reduced by 2-3 log_10_ as compared to untreated JEV-infected mice brains (Fig. 2G). One of the striking pathological features of JEV infection is encephalitis; hence we also tested the effect of TFP on virus induced neuroinflammation in these mice. TFP treatment subdued the production of pro-inflammatory cytokines (IFNB1, IL6, RANTES and TNFa) and protected the mice from JEV-induced neuroinflammation (Fig. 2H). Altogether these findings substantiate the antiviral efficacy of TFP against JEV in animal model.

### 3.3. Anti-DENV activity of TFP is compromised in IFNR deficient AG129 mice

TFP showed robust antiviral effect in Huh7 against DENV-2, with an IC50 value of 2.9 µM and CC50 value of 48.39 µM (Fig. 3A). A similar dose-dependent reduction in viral RNA was also observed (Fig. 3B). Its calculated SI was 16.7, making it a potential candidate to test in mice.

**Fig. 3:**
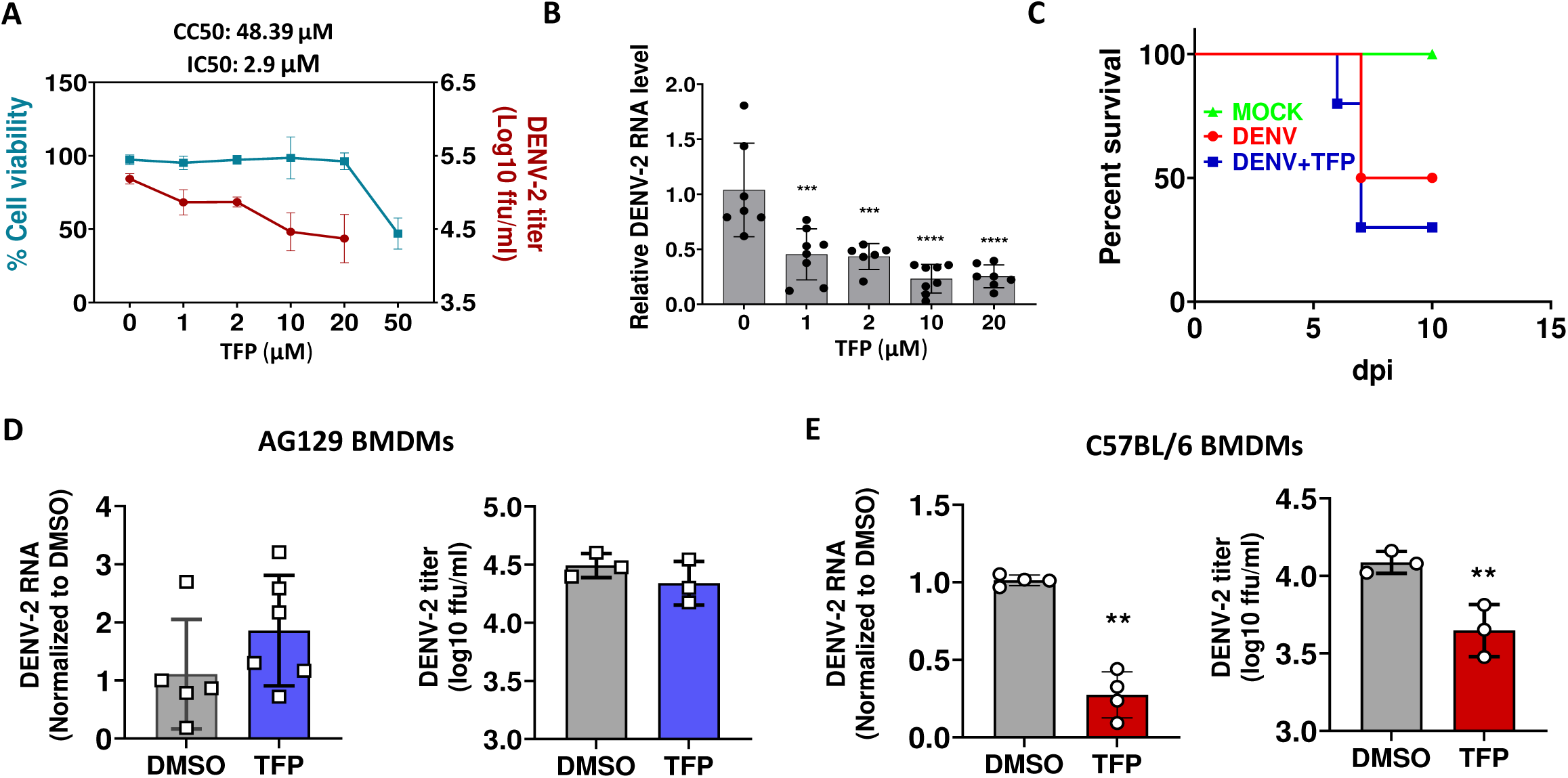
TFP exhibits antiviral activity against DENV-2 in Huh7 and C57BL/6 BMDMs, but loses its effect in the IFNR-deficient AG129 system. (A, B) Huh7 cells were mock/DENV-2 (2 MOI) infected for 48 h, followed by TFP treatment at indicated concentrations for 48 h. (A) DENV-2 titer was quantified in cell supernatant using foci-forming assay (*n*=6), and % cell viability was determined using MTT assay (*n*=4), as represented by line graphs. Means ± SD were plotted, and CC50 and IC50 values were calculated using non-linear regression analysis. (B) qRT-PCR was performed to determine intracellular DENV-2 RNA levels (*n*=6). One-way ANOVA with Dunnett’s test was used to determine the statistical significance of the data. (C) AG129 mice were intraperitoneally infected with 10^5 ffu of mouse-adapted DENV-2, followed by PEG400/TFP (1 mg/kg) oral administration at 4 hpi. The dosage was continued until day 8 post-injection (p.i.) at an interval of 24 h. Mice were observed till 10 dpi to obtain survival curves. Log-rank Mantel-Cox test was performed to define statistical significance. (D) BMDMs isolated from AG129 mice were infected with 5 MOI of DENV-2. Cells were either DMSO-treated or TFP-treated (10 µM) for 24 h. The intracellular viral RNA was quantified using qRT-PCR (left panel), and titers were calculated from cell supernatant using foci-forming assay (right panel). (E) C57BL/6 BMDMs were infected with DENV-2 (5 MOI) and DMSO/TFP (10 µM) treated till 24 hpi. Left panel represents intracellular DENV-2 RNA levels, and right panel represents DENV-2 titers. Viral RNA levels were normalized to the DMSO control. Values were plotted as *n*≥3. The unpaired Student’s t-test was used to calculate the *p* values: \**p*<0.05, \*\**p*<0.01, \*\*\**p*<0.001, \*\*\*\**p*<0.0001.

Till date, AG129 is the most sensitive model to study DENV infection *in vivo*. These mice lack both type 1 and type 2 interferon (IFN) response and hence support robust DENV replication. Therefore, in this study we used AG129 mice to examine *in vivo* efficacy of our drug of interest. Clinical signs such as pyrexia, piloerection and loss of body weight began to emerge by days 4-6 post infection. By the 7^th^ day, 50-70% of infected mice were dead, with no significant difference between the mice treated with vehicle and with drug (Fig. 3C). Median survival time calculated for untreated and TFP-treated cohorts of mice was 8.5 and 7 days, respectively. Consistent with the *in vivo* observation, TFP treatment did not exert any antiviral effect on AG129 derived BMDMs (Fig. 3D). However similar experiments performed in BMDMs isolated from C57BL/6 mice revealed that the drug can eliminate the virus in IFN receptor-competent WT background (Fig. 3E). Due to the lack of an appropriate WT mouse model for DENV, the *in vivo* effect of our drug against DENV-2 could not be studied. These data also highlight that functional IFN signaling is essential for the antiviral effect of TFP.

### 3.4. TFP demonstrates robust antiviral activity against CHIKV in cell line and mice model

As observed for the other two cell lines, ERMS did not show any discernible cytotoxic effects of TFP at concentration ≤ 20 µM. The CC50 (49.22 µM) of TFP in these cells was notably higher than IC50 (0.74 µM) of the drug (Fig. 4A), culminating a favorable SI of 66.5. Dose-response curve of intracellular CHIKV RNA also showed a significant drop in presence of TFP (Fig. 4B). Furthermore, TFP treatment strongly inhibited CHIKV replication in murine primary BMDMs (Fig. 4C). These findings suggest that the antiviral activity of TFP against CHIKV is consistent and robust.

**Fig. 4:**
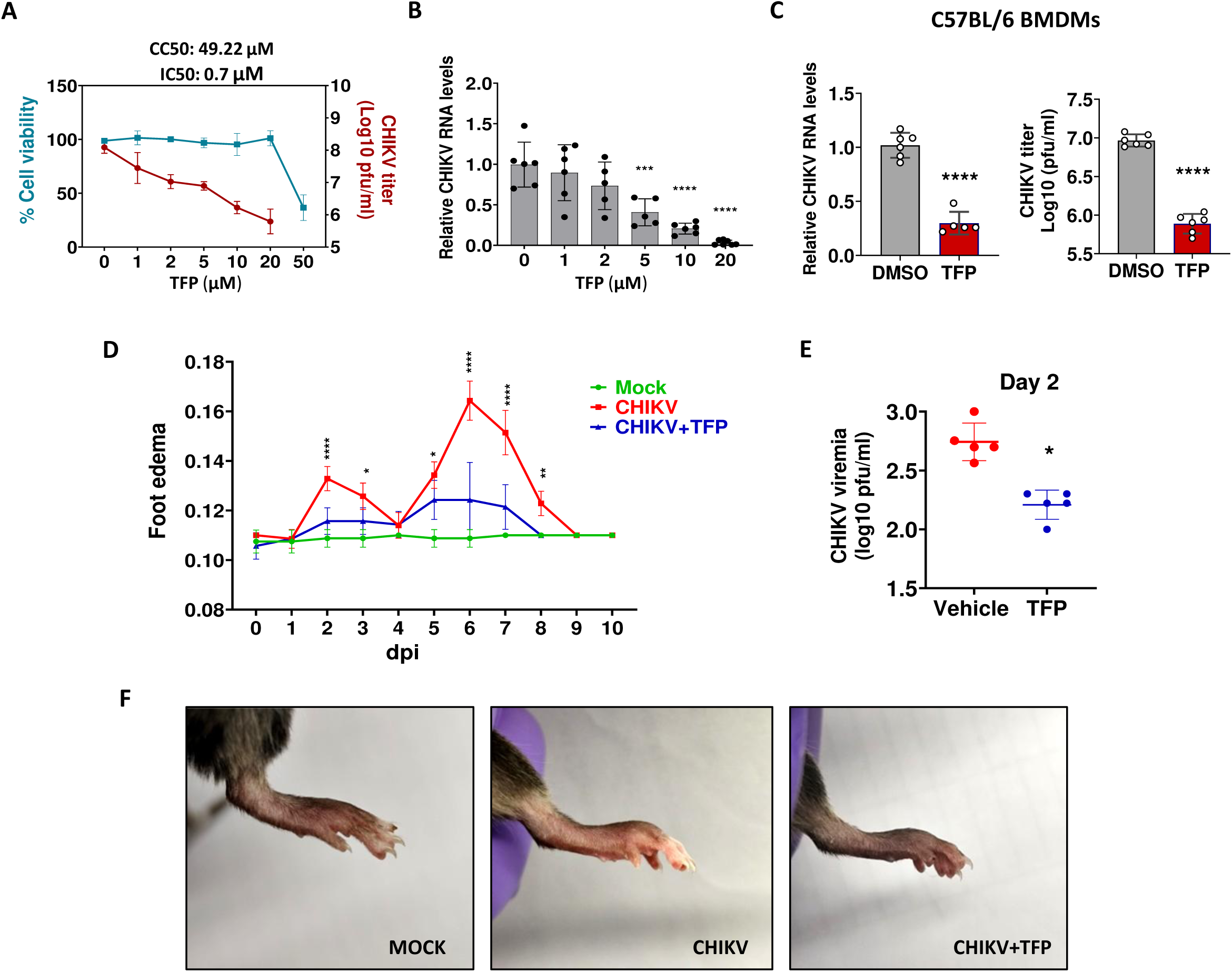
TFP restricts CHIKV infection in cell line and mice model. (A, B) ERMS cells were mock- or CHIKV-infected and treated with DMSO/TFP at the indicated concentrations until 24 hpi. (A) Percent cell viability (*n*=4) and viral titers (*n*=6) were plotted as mean ± SD. Non-linear regression analysis was used to calculate CC50 and IC50 values. (B) Total RNA was isolated from cells at 24 hpi, and qRT-PCR was performed to quantify CHIKV RNA levels. Data were normalized to DMSO control (*n*=6). Statistical significance was determined using one-way ANOVA followed by Dunnett’s test. (C) BMDMs isolated from C57BL/6 mice were infected with CHIKV at 1 MOI and treated with DMSO/TFP (10 µM) for 24 h. Left panel shows intracellular CHIKV RNA levels (*n*=6) normalized to DMSO, and right panel shows extracellular CHIKV titer (*n*=6). The unpaired Student’s t-test was used to determine the statistical significance. (D) Twelve-week-old C57BL/6 mice were mock-infected or infected with 10^4 pfu of CHIKV subcutaneously. Infected mice were treated with vehicle (PEG400) or TFP (1 mg/kg) orally. Treatment plan was same as used in JEV and DENV-2 infection models. All the mice were monitored until 10 dpi, and the paw edema was measured daily using a plethysmometer and plotted as line graphs. Two-way ANOVA test followed by Sidak’s multiple-comparison test was used to determine the statistical significance between CHIKV (*n*=6) and CHIKV+TFP treated (*n*=6) groups. (E) The viral load in serum of vehicle-treated (*n*=4) and TFP-treated (*n*=6) CHIKV-infected mice was determined at 2 dpi using plaque assay. The unpaired Student’s t-test was used to calculate statistical significance. (F) The representative images of paw swelling at 6 dpi, in different treatment groups, are shown here. \**p*<0.05, \*\**p*<0.01, \*\*\**p*<0.001, \*\*\*\**p*<0.0001.

To test the *in vivo* antiviral efficacy of our drug against CHIKV, we used sub-lethal mice model wherein the characteristic disease signs of CHIKV infection- fever and arthropathy- were observed [33]. Subcutaneous inoculation of CHIKV (10^5 pfu/mice) in hind footpad of C57BL/6 mice, led to development of transient edema and detectable viremia. In vehicle-treated CHIKV-infected mice, we observed biphasic inflammatory response where first small peak of paw edema began to appear at 2 dpi, self resolved in next 2 days, and further attained maximum peak at 6 dpi (Fig. 4D). Whereas mice treated with TFP (1 mg/kg, daily) showed minimum swelling (Fig. 4D). As compared to vehicle-treated mice, these mice exhibited a significant reduction in paw volumes across all time points. This difference in CHIKV-induced paw inflammation between vehicle-treated and drug-treated group can also be evidently seen through exemplar images obtained at 6 dpi (Fig. 4F).

The antiviral impact of TFP was further confirmed by quantifying the viral titer in serum of CHIKV-infected mice (Fig. 4E). We observed detectable viremia at 2 dpi, after which no infectious particle could be identified through plaque assay. It could be due to the systemic clearance of CHIKV. However, on day 2 we could observe that mice treated with TFP exhibited ∼70% reduction in serum viral load compared to control, establishing the robust antiviral potential of drug against CHIKV (Fig. 4E).

### 3.5. TFP induced ER stress is essential for its broad spectrum antiviral activity

Prior studies have reported that the anti-JEV activity of phenothiazines is mediated through the induction of ER stress [26, 27]. To investigate whether this mechanism corresponds to TFP’s antiviral mode of action, we began by quantifying the levels of eIF2α phosphorylation –a marker for ER stress induction. We found that TFP in Neuro2a cells shows robust increase in phosphorylation of eIF2a after 3 hours of treatment (Fig. 5A, left). The expression of total protein, here, is also enhanced. The phosphorylation and expression pattern of eIF2a in Thapsigargin (Tg) treated Neuro2a cells (positive control) was found to be identical with TFP treated cells, confirming that TFP is a potent ER stress inducer. By 6 hpt (hours post treatment), we continue to observe p-eIF2a induction in Tg- and TFP- treated Neuro2a cells. However, the extent of phosphorylation declined and the level of total protein returns to basal level (Fig. 5A, right). These phosphorylation dynamics can be explained by the classic phenomenon of integrated stress response where the activation of negative feedback regulators over time reset the ER stress towards baseline. These observations were also validated in Huh7 and ERMS cells. In accord with our findings from Neuro2a cell line, TFP induced the phosphorylation of eIF2a and enhanced its protein abundance at 3 hpt and 6 hpt in both the cell lines (Fig. 5C, E). A similar increase in p-eIF2a and eIF2a levels was also shown by Tg treatment. Altogether these results suggest that TFP induced ER stress is a consistent phenomenon detected across cell lines of different lineages.

**Fig. 5:**
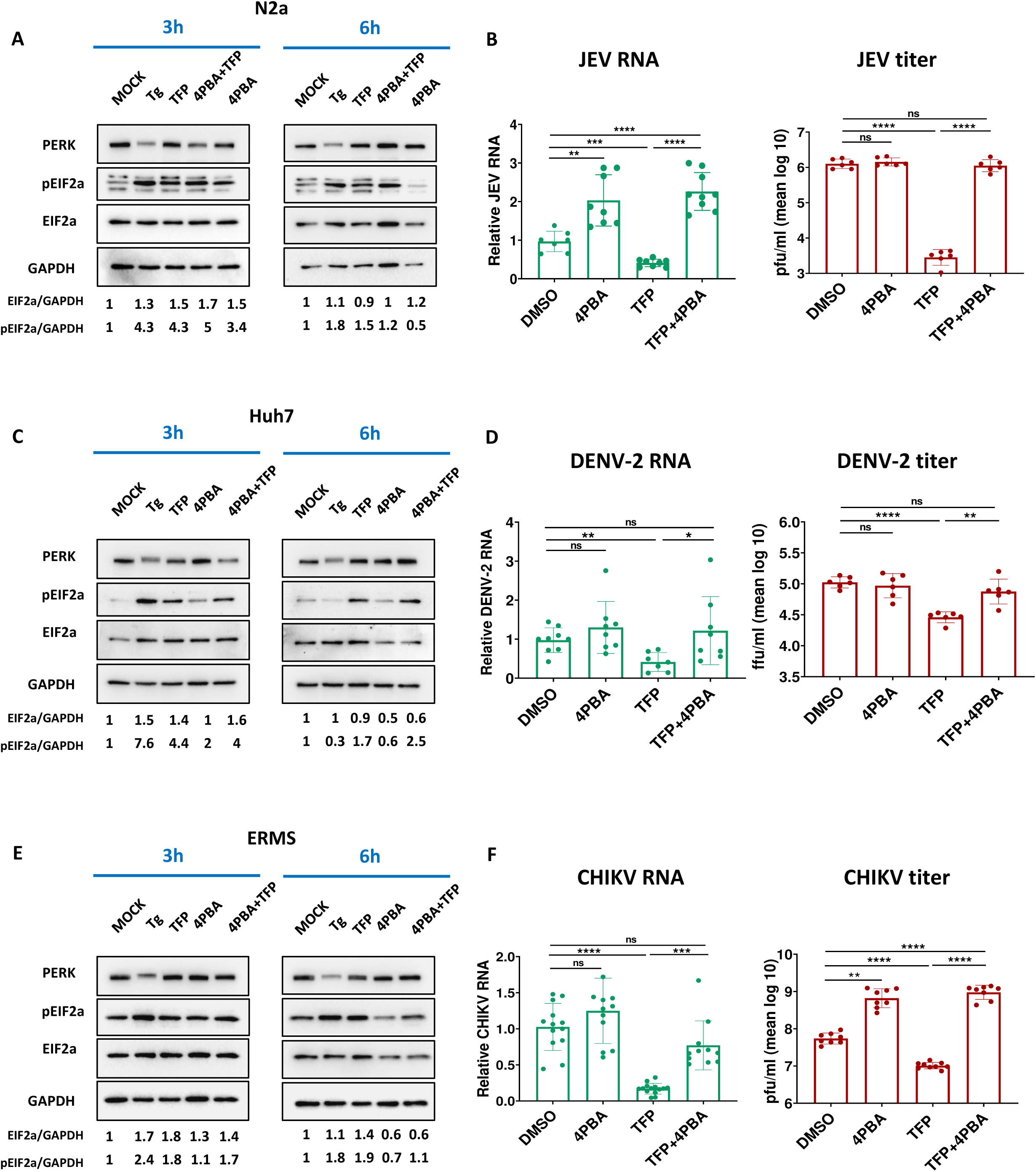
TFP-induced ER stress mediates its broad-spectrum antiviral activity. (A, C, E) Neuro2a (A), Huh7 (C), and ERMS (E) were treated with DMSO/Tg (1 µM) /TFP (10 µM)/4PBA (2 mM)/ TFP+4PBA for 3 h and 6 h. Cell lysates were prepared, and proteins were immunoblotted for p-eIF2a, eIF2a, PERK, and GAPDH (loading control). The values below the blot show relative expression of p-eIF2a/GAPDH and eIF2a/GAPDH after normalization to DMSO-treated cells. (B) Neuro2a cells were infected with JEV at 1 MOI, followed by treatment with DMSO/4PBA (2mM)/TFP (10 µM)/4PBA+TFP for 24 h. Left panel displays the quantified intracellular JEV RNA levels, and right panel shows JEV titers from two independent experiments (*n*=6). (D) Huh7 were infected with DENV at 2 MOI and treated with DMSO/4PBA (2mM)/TFP (10 µM)/4PBA+TFP for 48 h. Left panel shows the bar graph with intracellular viral RNA levels normalized to DMSO (*n*=6), and right panel shows extracellular virus titers calculated using foci-forming assays (*n*=6). (F) ERMS cells infected with CHIKV (1 MOI) and treated with DMSO/4PBA (2mM)/TFP (10 µM)/4PBA+TFP were compared to quantify CHIKV RNA (left panel) and CHIKV titer (right panel). RNA levels were normalized to DMSO (*n*=6), and titers were plotted as mean log10 pfu/ml (*n*=6). The unpaired Student’s t-test was used to determine the *p* values: \**p*<0.05, \*\**p*<0.01, \*\*\**p*<0.001, \*\*\*\**p*<0.0001.

Additionally, we observed that Tg treatment in all these cell types led to shift in mobility of PERK, indicating its phosphorylation. On the other hand, no such pattern was found in TFP treatment (Fig. 5A, B). Hence, we can conclude, in line with our previous report, the nature of ER stress induced by phenothiazine is adaptive and different from ER stress induced by Tg [26].

We also observed that treatment with 4PBA (4-phenylbutyric acid), a chemical chaperone, relieves ER stress and reduces the phosphorylation of eIF2a in a cell-agnostic manner (Fig. 5A, C, E). Furthermore, its co-treatment with TFP reduced the phosphorylated and total eIF2a levels, indicating the attenuation of TFP-induced ER stress.

To determine if the antiviral effect of TFP is attributable to ER stress induction, we assessed if 4PBA treatment could counteract the drug’s antiviral activity by restoring ER proteastasis. As we co-treated the cells, we observed complete rescue of virus replication (Fig. 5B, D, F). The antiviral effect of TFP on JEV RNA and titer was entirely abrogated upon treatment (Fig. 5B). 4PBA also rescued viral RNA levels and foci forming units of DENV-2 (Fig. 5D). Similar findings were obtained for CHIKV infections (Fig. 5F). Taken together these observations corroborate that TFP-induced ER stress is central to its broad-spectrum antiviral mode of action.

## 4. Discussion

Arboviral infections pose a major global threat to human health, causing hundreds of thousands of deaths annually. Despite the availability of vaccines, risk of emergence and re-emergence of arboviruses has increased in the past few decades. Many global initiatives are underway to address potential outbreaks, including implementation of effective surveillance systems, vector management programs, and identification of areas at risk of emergence (WHO Global Arbovirus Initiative, 2022). But these efforts seem to hit a roadblock in therapeutics. Due to a lack of specific antivirals, existing treatment is limited to supportive care. Hence, it becomes essential to develop broad-spectrum antivirals against arboviruses.

Amongst arbovirus-associated diseases, infections caused by DENV, CHIKV, ZIKV, YFW, JEV and WNV are most prevalent [34]. In this study, we have demonstrated broad-spectrum antiviral activity of trifluoperazine against three clinically relevant arboviruses: DENV, CHIKV, and JEV.

Trifluoperazine (TFP) is a widely prescribed first-generation antipsychotic drug. It is a phenothiazine derivative, extensively reported for its repurposing potential as anti-proliferative [23, 35–38], immunomodulatory [39–41], neuroprotective [42–45], anti-bacterial [46–48] and antiviral agent [25, 27, 49–52]. Most of these properties of TFP are attributable to its dual antagonism of dopamine receptor D2 (D2R) and calmodulin (CaM). As a D2R antagonist, TFP primarily exerts an inhibitory effect on dopaminergic signalling and offers neuroprotective benefits. Alternatively, it affects cell proliferation by increasing cAMP production, activating MAPK or AMPK, and reducing beta-catenin degradation [53]. By binding to calmodulin, TFP alters the intracellular calcium flux [54], which consequently modulates the downstream cellular processes, such as proliferation and differentiation pathways [55, 56], metabolic pathways [57, 58], and stress pathways [26, 27, 59]. One of such stress responses is ER stress. TFP binding to calmodulin leads to the activation of ER calcium channels and depletion of intraluminal calcium ions, resulting in the induction of ER stress and activation of the unfolded protein response [60–63].

Our study demonstrates that TFP treatment induces ER stress in diverse cell types, suggesting the phenomenon is not cell-specific. Through temporal expression profiling, we have shown an early and robust increase in phosphorylated eIF2α, which subsequently declines at later time points. It is likely because drug-induced ER stress activates the UPR, which attempts to restore cellular homeostasis. This reversible induction of ER stress describes the adaptive phase of the stress response and is used to target virus infections.

A large number of RNA viruses, including arboviruses, have been reported to trigger ER stress sensors [9]. ER stress is a fundamental host defence mechanism, and its pharmacological induction has been reported to exhibit antiviral activity against many viruses. For example, increasing the phosphorylated pool of eIF2a by small molecules has been shown to inhibit the replication of KSHV, HSV and DENV [64–66], targeting the IRE1 and XBP1 pathway has shown broad-spectrum antiviral activity against SINV, MNV-1 and La Crosse Virus [21], and acute induction of ER stress by tunicamycin or thapsigargin has been shown to limit the infection of SARS-CoV-2 [67]. Also, our previous study has demonstrated that ER stress-induced autophagy by methotrimeprazine restricts JEV infection [26]. In this study, we have shown that TFP-induced adaptive ER stress is central to its broad-spectrum antiviral activity against arboviruses. Since all the tested viruses depend on ER for their replication and translation, early induction of ER stress by TFP made the cellular environment hostile and less permissive for replication. We have also demonstrated that treatment with the chemical chaperone 4PBA alleviated the TFP-induced ER stress, rendering infected cells once more susceptible to sustaining infection.

We observed high selectivity indices for antiviral activity of TFP against JEV, DENV-2 and CHIKV in cell lines. It prompted us to validate the drug in respective mouse models. We observed that, in JEV-infected C57BL/6 mice, TFP treatment significantly reduced the viral load and associated pathogenesis, thereby providing survival benefits to the mice. Owing to the drug’s ability to cross the BBB, it is possible that TFP restricts the JEV infection primarily in the central nervous system (CNS). However, we cannot rule out the possibility of peripheral clearance of the virus for two key reasons. First, we observed antiviral activity of TFP in peripheral immune cells, BMDMs. Second, the anti-JEV activity of TFP is ER stress-dependent, and many cell types present in the periphery are capable of inducing ER stress.

Being efficiently armed with an IFN response, C57BL/6 mice rapidly clear DENV; hence, we used IFN receptor knockout (IFNR-/-) AG129 mice to test TFP. Surprisingly, the antiviral effect of TFP against DENV-2 was lost in AG129 mice. However, comparing the antiviral activity of TFP against DENV-2 in BMDMs isolated from WT C57BL/6 mice and AG129 mice revealed that the drug can eliminate the virus in the WT background. Therefore, we speculated that the antiviral effect of TFP against DENV-2, in addition to being ER stress-dependent, is attributable to a functional IFN response. One plausible explanation is that TFP-induced phosphorylation of eIF2α promotes the formation of stress granules, which in turn sequester viral RNA/proteins for their elimination by PRRs and associated interferon response [68, 69]. Nevertheless, this hypothesis warrants further exploration. It brings us to the limitation of this study, which is the lack of an IFNR-competent mouse model to test our drug against DENV infections. However, the repurposing potential of TFP against DENV-2 cannot be undermined solely because it fails in the AG129 mouse model. AG129 mice exhibit systemic features similar to those seen in dengue patients, but they do not replicate the functional immune interactions necessary to mimic actual DENV infection in humans.

TFP also showed promising results in its preclinical testing against CHIKV infection. As evidenced by reduced viremia and attenuated paw inflammation, TFP treatment resulted in rapid clearance of circulating and disseminated virus. The observed broad-spectrum *in vivo* antiviral efficacy of the drug strengthens our confidence to develop TFP as a clinical candidate. It is further supported by well-documented pharmacokinetic and pharmacodynamic profiles of the drug, which can alleviate the challenges of early-phase clinical trials, thus accelerating the timeline of its clinical translation. Moreover, the host-centric antiviral mode of action of the drug reduces the likelihood of rapid virus resistance, which is a concern associated with the translation of direct-acting antivirals.

## Acknowledgements

We acknowledge the facilities and staff of Experimental Animal Facility (EAF) of NCR Biotech Science Cluster, Faridabad. We also thank Prof. Prasanjit Guchhait, for mouse adapted DENV-2 virus strain (P8-P23085 INDI-60), and Prof. Sudhanshu Vrati for CHIKV INDI-06-Guj isolate. We extend our acknowledgement to Mr. Shouri KA and Ms. Khashpatika Ganesh for their help in conducting CHIKV animal experiments.

